# Deletion or Targeted Blockade of FcγRIIb (CD32b) Impairs α-Syn Propagation *In-Vivo*

**DOI:** 10.1101/2025.11.18.687143

**Authors:** James M. Hennegan, Michael J. Hurley, Kerry L. Cox, Leon R. Douglas, Patrick J. Duriez, Robert J. Oldham, Mark S. Cragg, Björn Frendéus, Ali Roghanian, Jessica L. Teeling

## Abstract

Parkinson’s disease (PD), the most common neurodegenerative movement disorder, is characterised by pervasive deposition of alpha-synuclein (α-Syn) aggregates and the death of dopaminergic neurons. Fc gamma receptor IIb (FcγRIIb or CD32b), the sole inhibitory FcγR in humans (h) and mice (m), serves as a molecular conduit for intercellular α-Syn transmission *in-vitro*. Here, we demonstrate that FcγRIIb facilitates α-Syn propagation and neurotoxicity *in-vivo* using the pre-formed fibril (PFF) α-Syn model in mice. Genetic ablation of mFcγRII attenuated PFF α-Syn-induced Lewy pathology, dampened associated neuroinflammatory responses, and preserved nigrostriatal dopaminergic neurons. Furthermore, pharmacological blockade of hFcγRIIb using clinically-related monoclonal antibodies mitigated acute-phase α-Syn pathology in hFcγRIIb-transgenic mice following PFF challenge. Collectively, our results indicate that FcγRIIb is a mediator of α-Syn propagation *in-vivo* and highlight it as a tractable therapeutic target against α-synucleinopathies like PD.

## Introduction

Central to Parkinson’s disease (PD) pathophysiology is alpha-synuclein (α-Syn), an intrinsically unstructured, presynaptically enriched protein that is prone to misfolding and assembly into β-sheet-rich conformers [1, 2]. Aggregated α-Syn drives the formation of signature Lewy bodies (LB) and Lewy neurites, which spread along interconnected brain circuitry in a predictable spatiotemporal pattern [3]. Prevailing evidence supports a prion-like mechanism of α-Syn propagation, whereby fibrillar seeds of misfolded α-Syn disseminate trans-neuronally, templating the aggregation of naïve monomeric (Mono) α-Syn into toxic oligomers or fibrils [4–7]. However, despite extensive characterisation of α-Syn aggregation and spread, the molecular mediators that govern the propagation of α-Syn remain incompletely defined. Multiple non-mutually exclusive mechanisms have been proposed, including tunnelling nanotube-mediated transfer, passive trans-membrane diffusion, and receptor-dependent endocytic uptake [8, 9].

Fc gamma receptor IIb (FcγRIIb or CD32b), referred to as FcγRII or CD32 in mice, has recently emerged as a promising receptor candidate implicated in α-Syn propagation, based on its capacity to bind aggregated α-Syn species [10, 11]. FcγRs comprise six isoforms in humans and four in mice [12–14], collectively orchestrating peripheral immune effector responses through a balance of activating and inhibitory signalling. Their principal role is to recognise and bind the Fc domain of IgG antibodies, coupling antigen recognition with downstream cellular responses such as phagocytosis and cytokine release, contributing to the preservation of peripheral immune homeostasis [12]. FcγRIIb is the sole inhibitory FcγR, conserved between species, and uniquely restrains immune cell activation following immune complex engagement, in part, *via* its immunoreceptor tyrosine-based inhibitory motif (ITIM) [15, 16]. Importantly, FcγRIIb expression is documented on neuronal and microglial membranes within the central nervous system (CNS), particularly under conditions of proteopathic stress [17–20].

Recently, Choi *et al*. provided direct *in-vitro* evidence that fibrillar species of α-Syn bind FcγRIIb to activate downstream SHP-1/2-c-Src signalling [10, 11, 21]. Neuronal FcγRIIb facilitates the internalisation and neurotoxicity of amyloidogenic proteins, including α-Syn and β-amyloid aggregates [11, 20]. In parallel, engagement of microglial FcγRIIb by α-Syn fibrils *in-vitro* suppresses phagocytic activity and by extension, clearance of extracellular debris like α-Syn, suggesting a dual role for FcγRIIb [10]. Interestingly, perturbation of the FcγRIIb-ITIM-SHP-1/2-c-Src axis *via* gene knockdown or pharmacological inhibition diminished α-Syn uptake and its intercellular transmission [11, 21]. Together, this suggests that FcγRIIb signalling may be a critical mechanism in α-Syn propagation, although the functional relevance of FcγRIIb *in-vivo* remains unresolved.

Here we used the pre-formed fibril (PFF) α-Syn model to investigate the role of mFcγRII towards α-Syn propagation *in-vivo* using wild-type (WT) and murine FcγRII-deficient (mFcγRII^-/-^) C57BL/6 mice [22]. The translational potential of targeting human FcγRIIb (hFcγRIIb) was also explored using clinically-related monoclonal antibodies (mAb) displaying either Fc-WT, or N297Q mutant (Fc-Null) Fc domains, in transgenic mice expressing hFcγRIIb (hFcγRIIb Tg) [23–25]. Both mAb formats were designed to antagonise hFcγRIIb selectively, inhibiting ITIM activation and downstream signalling [23]. Herein, we report that both genetic knockout of mFcγRII and prophylactic hFcγRIIb blockade successfully mitigate α-Syn propagation in the PFF α-Syn mouse model.

## Results

### Genetic deletion of *Fcgr2b* eliminates mFcγRII expression in the adult mouse brain

To establish a validated foundation for investigating the role of mFcγRII in α-Syn propagation within the brain, we first sought to verify previous reports of mFcγRII expression in the adult mouse brain [17, 18, 20]. We evaluated mFcγRII expression in the brain of WT and homozygous mFcγRII^-/-^ mice; the latter originally generated *via* targeted deletion of exons 4 and 5 of the mouse *Fcgr2b* gene [22], resulting in a functional receptor knockout model. Examination of *in-situ* hybridisation data from the Allen Mouse Brain Atlas (https://mouse.brain-map.org/) revealed diffuse yet discernible *Fcgr2b* mRNA signals throughout the WT C57BL/6 brain (Fig. S1). Consistent with these observations, RT-PCR analysis detected robust *Fcgr2b* transcript levels in WT brain homogenates, comparable to those in spleen - a tissue with high mFcγRII abundance [24]. Conversely, *Fcgr2b* transcripts were undetected in both brain and spleen homogenates from mFcγRII^-/-^ mice (Fig. 1a). Importantly, expression of other FcγR isoforms was unaffected by *Fcgr2b* deletion, confirming the selectivity of the knockout (Fig. 1a). At the protein level, immunodetection of mFcγRII in WT tissue sections demonstrated reproducible labelling across multiple brain regions, including the Substantia Nigra *pars compacta* (SNpc; Fig. 1b), and confirmed in the spleen (Fig. 1c). No protein signal was observed in corresponding regions of mFcγRII^-/-^ mice (Fig. 1b, c). These data confirm that mFcγRII is expressed across the adult WT mouse brain, and its expression is abolished in mFcγRII^-/-^ mice, establishing a robust genetic framework for functional studies of mFcγRII.

**Figure 1.**
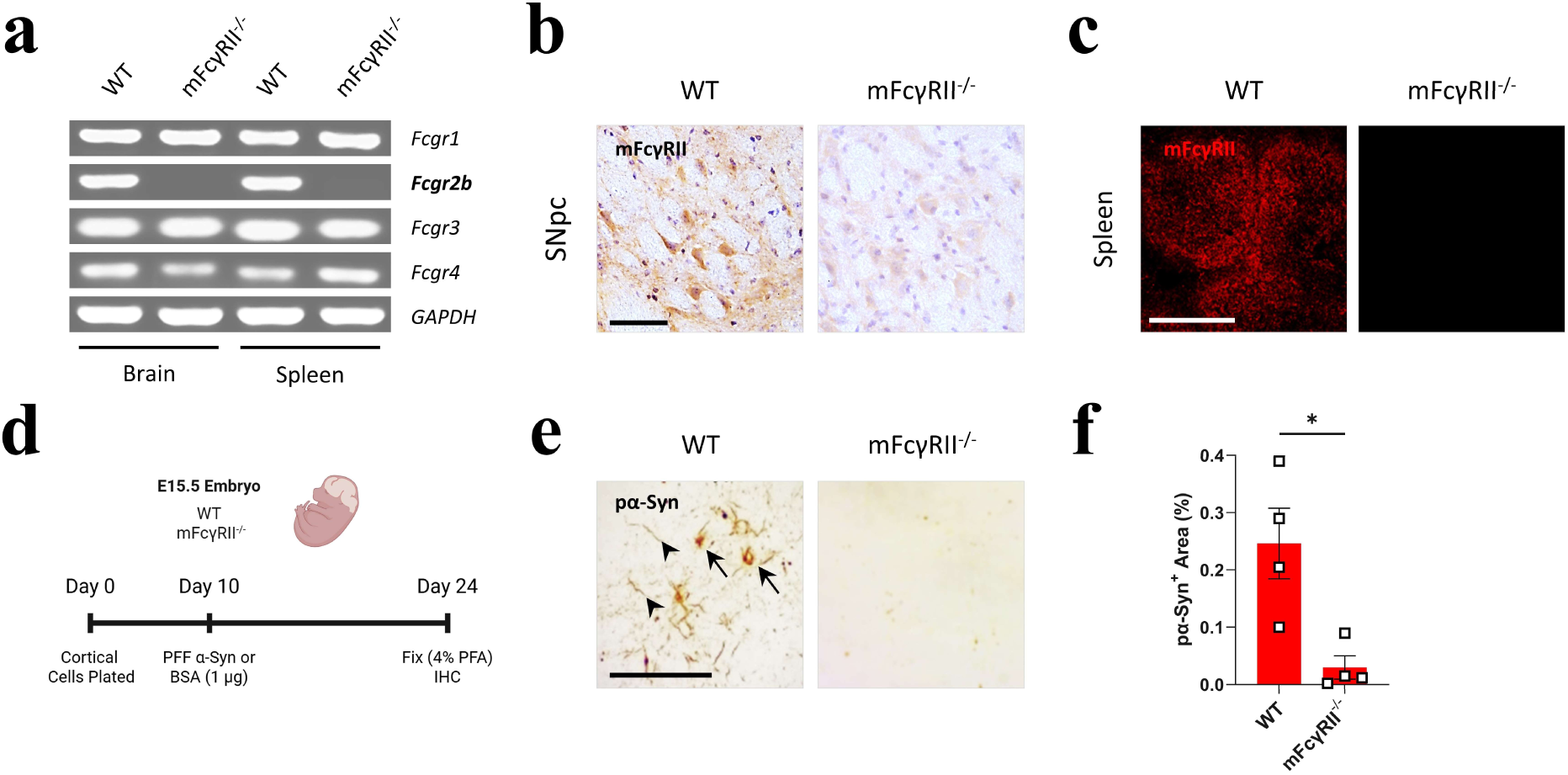
*Fcgr2b* deletion reduces pα-Syn accumulation in primary neurons from mFcγRII^-/-^ mice. **(a)** Agarose gel showing FcγR transcript expression in naïve mouse tissues. Total RNA from brain and spleen of WT and mFcγRII^-/-^ mice was analysed by RT-PCR using primers for mFcγRI (*Fcgr1*), mFcγRII (*Fcgr2b*), mFcγRIII (*Fcgr3*), mFcγRIV (*Fcgr4*), and *GAPDH* as house-keeping control. **(b)** Representative images of immunohistochemical mFcγRII staining in the Substantia Nigra *pars compacta* (SNpc) from WT and mFcγRII^-/-^ mice. Sections were counterstained with haematoxylin. Scale bar = 50 μm. **(c)** Representative images of immunofluorescent mFcγRII staining (red) in the spleen of WT and mFcγRII^-/-^ mice. Scale bar = 200 μm. **(d)** Schematic representation of *in-vitro* primary neuronal culture experiment. **(e)** Representative images of immunocytochemical pα-Syn^+^ staining in WT and mFcγRII^-/-^ primary neuronal cultures following 14 day PFF α-Syn (1 μg) incubation. Arrows denote pα-Syn localisation in perikarya. Arrowheads highlight neuritic structures. Scale bar = 100 μm. **(f)** Quantification of pα-Syn^+^ immunoreactivity (% area) in WT and mFcγRII^-/-^ cultures from **(e)**. Data are mean ± SEM from 4 separate experiments performed in triplicate. Statistical significance was determined using the Mann-Whitney test (\**p*<0.05).

### Phosphorylated (p) α-Syn is reduced in primary neurons from mFcγRII^-/-^ mice

To complement previous *in-vitro* studies using neuronal-like SH-SY5Y cells [11], and evaluate the role of mFcγRII in a physiologically relevant *ex-vivo* system, we generated and applied human PFF α-Syn (Fig. S2) to primary cortical neurons derived from E15.5 WT and mFcγRII^-/-^ embryos. Neurons were exposed to PFF at day 10, and the formation of pathological inclusions was assessed 14 days later (Fig. 1d) *via* immunostaining for α-Syn phosphorylated at serine 129 (pα-Syn) - the predominant pathological modification of α-Syn [26]. Neuronal cultures from WT mice developed abundant pα-Syn-positive (pα-Syn^+^) immunoreactivity, localising to both perikarya and neuritic arbours (Fig. 1e). Under identical conditions, mFcγRII^-/-^ neurons exhibited strongly reduced (88% less, *n* = 4) pα-Syn⁺ staining as compared to WT cultures (Fig. 1e, f). No pα-Syn signal was detected in control cultures treated with bovine serum albumin, confirming the specificity of the PFF-induced pα-Syn amplification (data not shown).

### Loss of mFcγRII attenuates seeding and propagation of α-Syn *in-vivo*

To determine the impact of mFcγRII deletion on the initiation and propagation of α-Syn pathology *in-vivo*, we used the PFF α-Syn model of synucleinopathy in mice, which recapitulates cardinal features of α-Syn seeding and LB expansion [27, 28]. Male WT and mFcγRII^-/-^ mice (2 months-old) received unilateral stereotaxic injections of human PFF α-Syn (*n* = 4, Fig. S2) or non-fibrillar control Mono α-Syn (*n* = 3) into the dorsal striatum. Mice were sacrificed and processed for immunohistochemical analyses at either 30 or 90 days post-injection (dpi) to capture emergent and developed stages of pathology, respectively (Fig. 2a).

**Figure 2.**
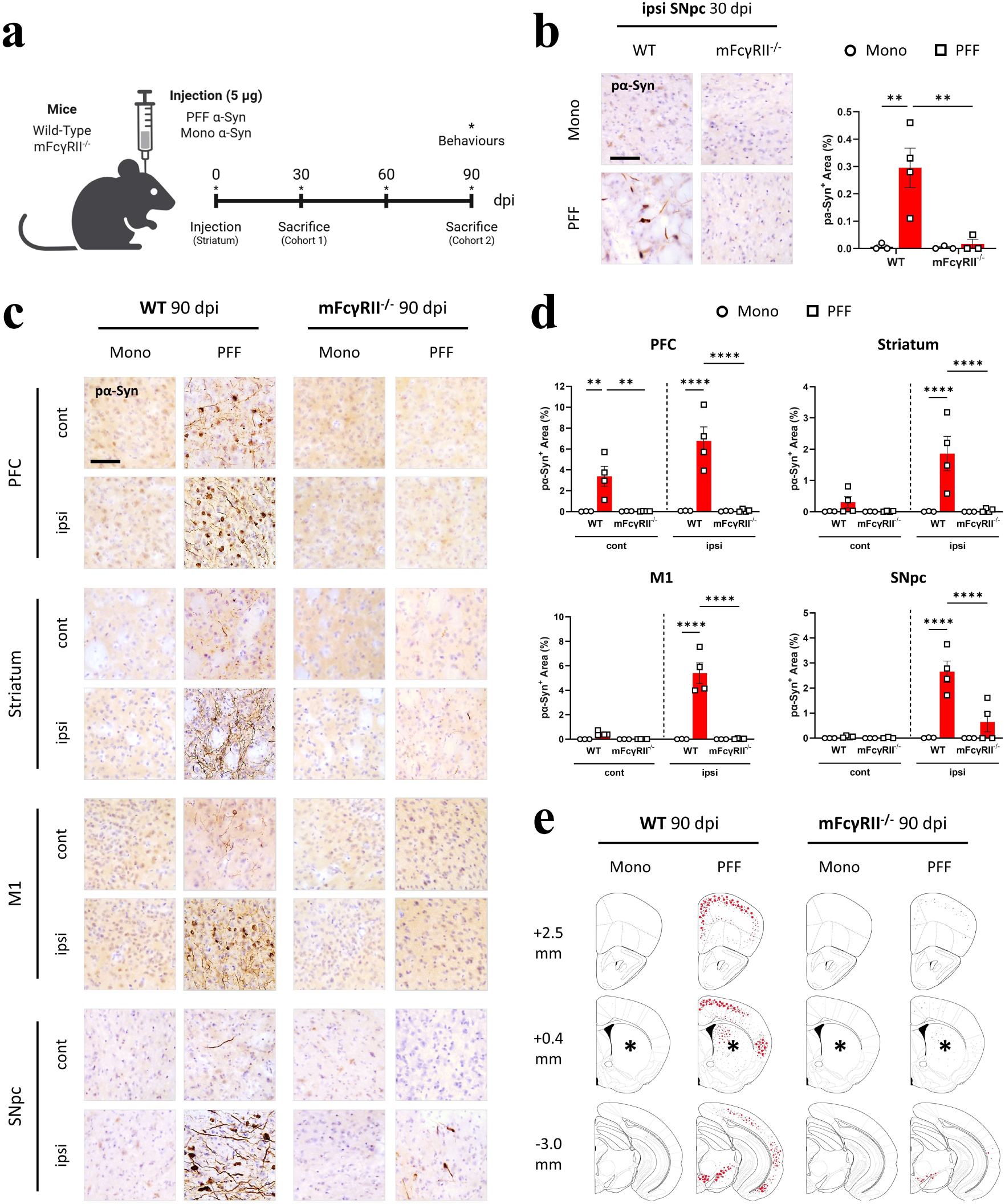
mFcγRII^-/-^ mice are protected from pα-Syn accumulation and anatomical distribution. **(a)** Schematic representation of *in-vivo* study design. **(b)** Representative images of immunohistochemical pα-Syn staining in the ipsilateral (ipsi) Substantia Nigra *pars compacta* (SNpc) of WT or mFcγRII^-/-^ mice injected with PFF or Mono α-Syn at 30 dpi. Sections were counterstained with haematoxylin. Scale bar = 50 μm. Graph shows quantification of pα-Syn^+^ immunoreactivity (% area) in indicated groups. **(c)** Representative images of immunohistochemical pα-Syn staining in contralateral (cont) and ipsi hemispheres of WT or mFcγRII^-/-^ mice injected with PFF or Mono α-Syn at 90 dpi. PFC; prefrontal cortex, M1; primary motor cortex. Sections were counterstained with haematoxylin. Scale bar = 50 μm. **(d)** Quantification of pα-Syn^+^ immunoreactivity (% area) at 90 dpi in indicated brain regions and groups from **(c)**. WT Mono; *n* = 3, mFcγRII^-/-^ Mono; *n* = 3, WT PFF; *n* = 4, mFcγRII^-/-^ PFF; *n* = 4 animals. Data are mean ± SEM. Statistical significance was determined using two-way ANOVA with Tukey’s multiple comparisons test (\*\**p*<0.01, \*\*\*\**p*<0.0001). **(e)** Distribution map of pα-Syn^+^ immunoreactivity (represented as red dots) at 90 dpi in the ipsi hemisphere of indicated groups. Asterisks denote the injection site (ipsi striatum), and adjacent numbers represent distance from Bregma in the anterior-posterior plane in mm.

In WT mice, PFF injection induced robust pα-Syn^+^ LB-like pathology as early as 30 dpi, localised to the ipsilateral (ipsi; side of injection) SNpc. In contrast, mFcγRII^-/-^ mice exhibited a near-complete absence of pα-Syn immunostaining at 30 dpi (Fig. 2b). Representative images of all regions of interest analysed at 30 dpi are presented in Figure S3a, revealing minimal pα-Syn^+^ immunoreactivity outside of the SNpc across all experimental groups (Fig. S3b). Notably, no pα-Syn^+^ signal was observed in WT or mFcγRII^-/-^ mice injected with Mono α-Syn at 30 dpi (Fig. S3a) or 90 dpi (Fig. 2c, Fig. S4a), confirming the specificity of the PFF-seeding response [27]. By 90 dpi of PFF, pα-Syn^+^ inclusions and neurites were abundant and widely distributed in the WT mouse brain, localising predominantly to the ipsi hemisphere. Brain regions with high pα-Syn^+^ immunoreactivity included the ipsi prefrontal cortex (PFC), injection site (striatum), primary motor cortex (M1), SNpc (Fig. 2c), entorhinal cortex (EC), and amygdala (Amy) (Fig. S4a). Analogous contralateral (cont) regions also exhibited muted pα-Syn deposition at 90 dpi (Fig. 2c, Fig. S4a), consistent with reported examples of trans-hemispheric propagation in the unilateral PFF injection model [27, 29].

Remarkably, mFcγRII^-/-^ mice injected with PFF exhibited sustained and profound suppression of pα-Syn^+^ immunostaining across all examined regions at 90 dpi. Sparse pα-Syn^+^ inclusions emerged in a subset of mice in select regions like the ipsi SNpc and Amy, although the overall pathological burden and its anatomical distribution were nominal as compared to WT mice (Fig. 2c, Fig. S4a). Quantitative image analysis revealed a significant reduction in total pα-Syn^+^ area (% coverage) across all examined brain regions in mFcγRII^-/-^ mice compared to WT mice (Fig. 2d, Fig. S4b). Reductions ranged from greater than 70% in the ipsi SNpc to almost total abrogation (∼99%) in the ipsi PFC and M1 (Fig. 2d), indicating impaired propagation of α-Syn along synaptically connected pathways. Figure 2e provides a regional distribution map of pα-Syn^+^ pathology throughout the ipsi hemisphere at 90 dpi, summarising the divergent pα-Syn expression patterns between WT and mFcγRII^-/-^ mice (expanded upon in Fig. S4c). Collectively, these data demonstrate that mFcγRII is not merely permissive but fundamental for efficient seeding and subsequent propagation of α-Syn pathology in the PFF model.

### Microglial reactivity in response to PFF α-Syn is attenuated in mFcγRII^-/-^ mice

Neuroinflammation is a central component of PD pathogenesis and a prominent feature of experimental mouse models of PD [30]. To assess whether deletion of mFcγRII modulates the neuroimmune response to PFF, we performed immunofluorescent labelling for Iba1, a pan-microglial marker, and FcγRI, the high-affinity IgG receptor that is increased on activated microglia [31]. In WT mice injected with PFF, we observed a pronounced elevation in microglial density within the ipsi striatum at 90 dpi, reflected by increased Iba1⁺ somatic staining (Fig. 3a, b). FcγRI immunoreactivity was likewise elevated at both 30 (Fig. S5a, c) and 90 dpi (Fig. 3a, c), indicating persistent microglial activation. Quantitative analysis of additional ipsi brain regions revealed sustained Iba1^+^ microglial density and FcγRI expression throughout the hemisphere at 90 dpi (Fig. 3d, e), aligning with the spatial distribution of pα-Syn, particularly in cortical regions.

**Figure 3.**
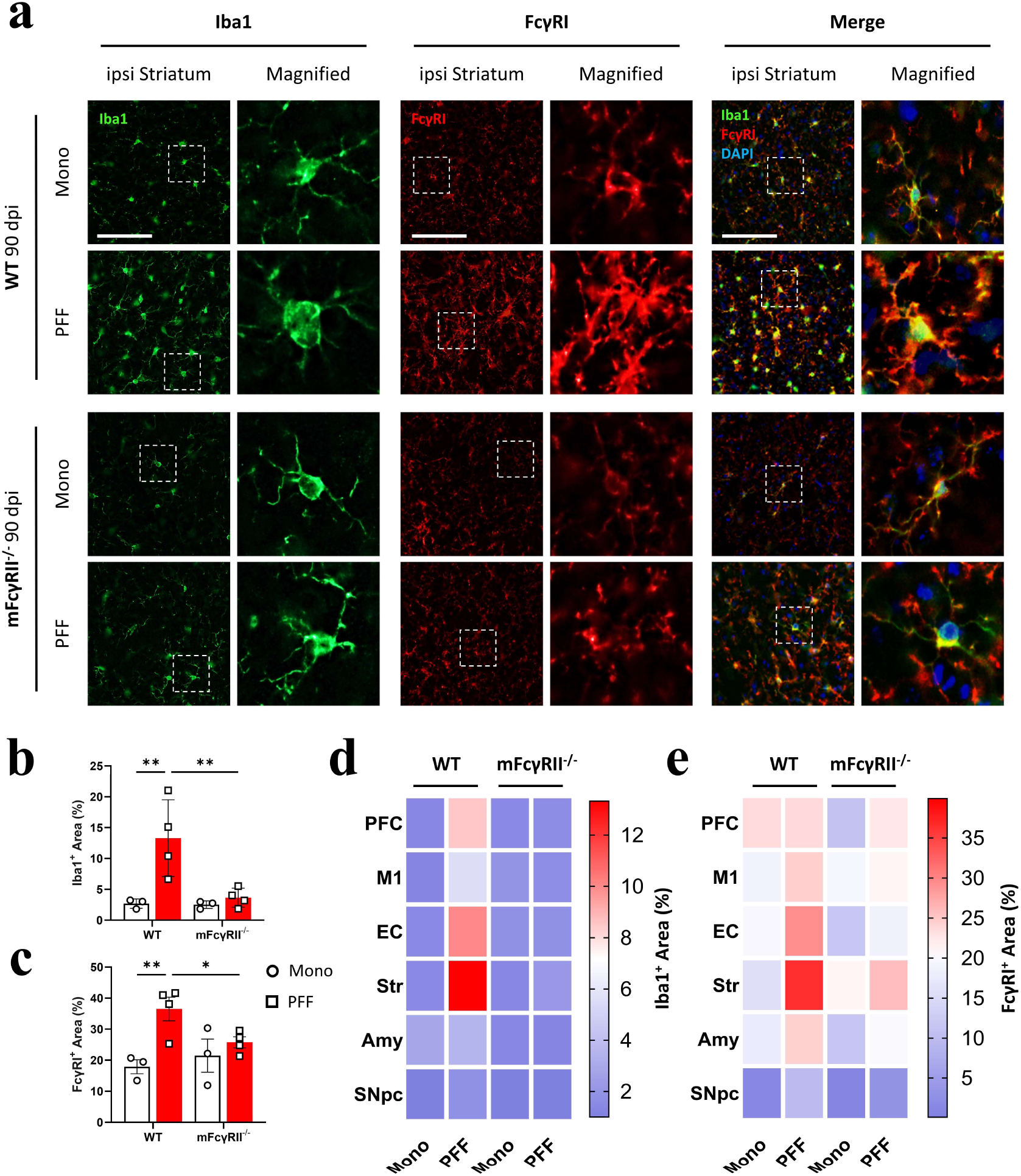
Microglial reactivity is reduced in response to PFF in mFcγRII^-/-^ mice at 90 dpi. **(a)** Representative images of immunofluorescent Iba1 (green), FcγRI (red), and co-staining (merge) in the ipsilateral (ipsi) striatum of WT and mFcγRII^-/-^ mice injected with PFF or Mono α-Syn at 90 dpi. Nuclei are labelled with DAPI (blue). Magnified images are shown in the right column of each panel. Scale bars = 50 μm. **(b, c)** Quantification of **(b)** Iba1^+^ and **(C)** FcγRI^+^ immunoreactivity (% area) in the ipsi striatum at 90 dpi. WT Mono; *n* = 3, mFcγRII^-/-^ Mono; *n* = 3, WT PFF; *n* = 4, mFcγRII^-/-^ PFF; *n* = 4 animals. Data are mean ± SEM. Statistical significance was determined using two-way ANOVA with Tukey’s multiple comparisons test (\**p*<0.05, \*\**p*<0.01). **(d, e)** Regional quantification of **(d)** Iba1^+^ and **(e)** FcγRI^+^ immunofluorescent staining at 90 dpi in different brain regions of interest. Each individual brain region is represented as a heat-map according to the mean immunopositive area (%) of each experimental group. PFC; prefrontal cortex, M1; primary motor cortex, EC; entorhinal cortex, Str; striatum, Amy; amygdala, SNpc; Substantia Nigra *pars compacta*.

By contrast, mFcγRII^-/-^ mice exhibited a substantially blunted microglial response following injection with PFF. At 90 dpi, we observed a ∼73% reduction in Iba1⁺ microglial density and a ∼29% decrease in FcγRI expression within the ipsi striatum relative to WT counterparts (Fig. 3b, c). A broader analysis revealed a comparatively diminished Iba1 and FcγRI immunoreactivity across the ipsi hemisphere of mFcγRII^-/-^ mice (Fig. 3d, e). Importantly, no significant microglial responses were observed in mice injected with Mono α-Syn (Fig. 3a-e). Together, these findings support the interpretation that microglial reactivity in our model correlates with regional pα-Syn deposition, the burden of which was substantially reduced in the mFcγRII^-/-^ brain.

### Motor function and nigrostriatal integrity are preserved in mFcγRII^-/-^ mice

The PFF model is well established for recapitulating progressive neurodegeneration, with selective vulnerability of dopaminergic neurons in the SNpc leading to behavioural deficits starting around 90 dpi [27]. Behavioural assays conducted prior to sacrifice at 90 dpi revealed that WT mice injected with PFF exhibited significant motor coordination impairments, evidenced by prolonged descent latency in the pole test (Fig. 4a). They also showed reduced locomotor activity and exploratory drive, presenting as diminished ambulation and spontaneous rearing in the open field test (Fig. 4b). On the contrary, mFcγRII^-/-^ mice maintained normal performance across all behavioural paradigms (Fig. 4a, b, S6), performing equivalently to Mono α-Syn-injected mice.

**Figure 4.**
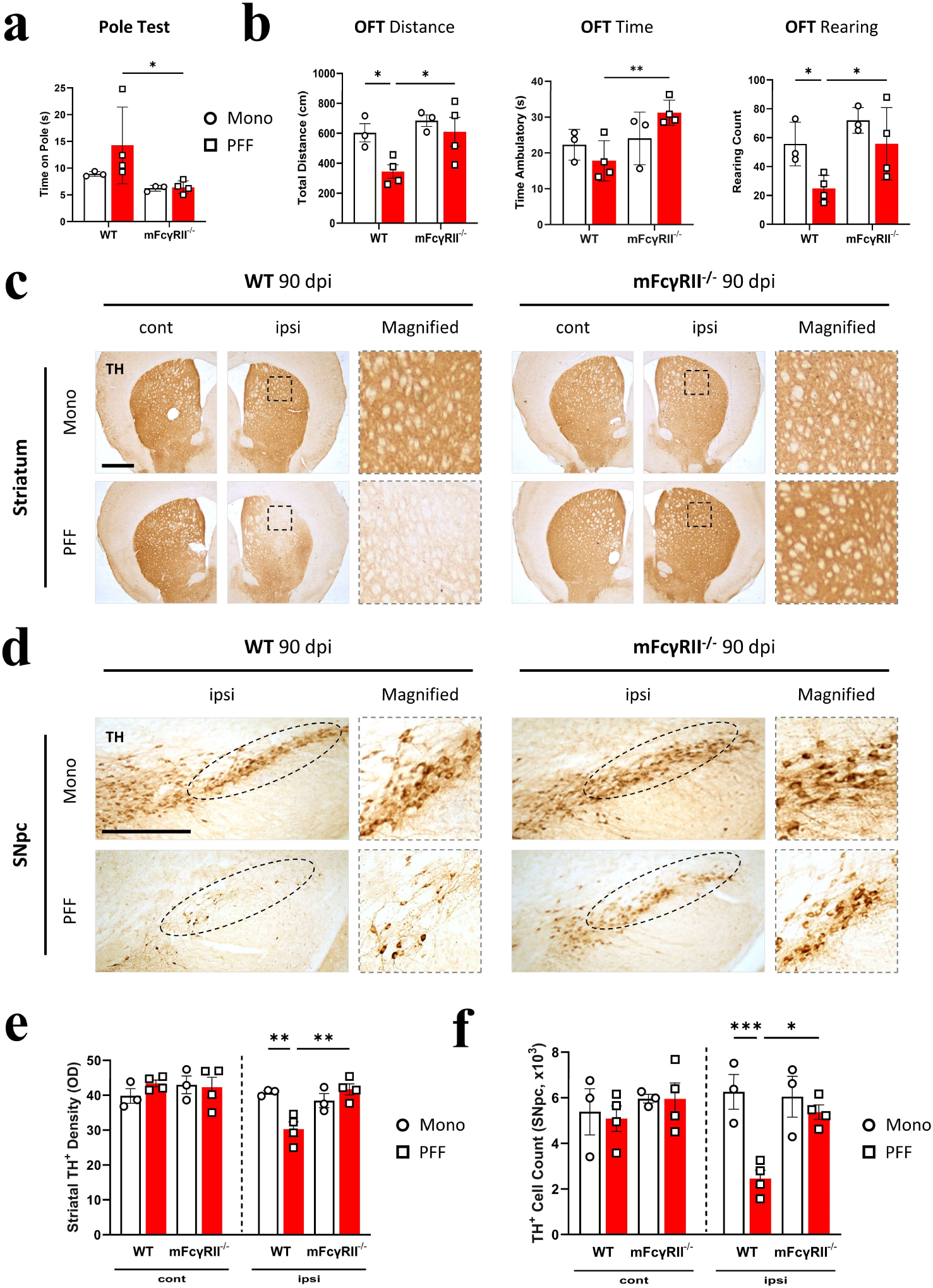
Genetic deletion of mFcγRII rescues motor dysfunction and dopaminergic neuronal integrity in response to PFF at 90 dpi. **(a)** Total time spent descending the pole in the pole test at 90 dpi. Each point represents the mean of 3 independent tests per mouse. **(b)** Total distance travelled, time ambulatory, and rearing count in the open-field (OFT) test at 90 dpi. **(c, d)** Representative images of immunohistochemical TH staining in **(c)** the contralateral (cont) and ipsilateral (ipsi) striatum and **(d)** the ipsi Substantia Nigra *pars compacta* (SNpc) of WT and mFcγRII^-/-^ mice injected with PFF or Mono α-Syn at 90 dpi. Magnified images are shown adjacent to each panel. Scale bars = 1 mm **(c)** and 200 μm **(d)**. **(e)** Quantification of TH^+^ optical density (OD) in the dorsal striatum and **(f)** TH^+^ cells in the SNpc represented as a number normalised to the total SNpc volume *via* the Cavalieri estimate method. WT Mono; *n* = 3, mFcγRII^-/-^ Mono; *n* = 3, WT PFF; *n* = 4, mFcγRII^-/-^ PFF; *n* = 4 animals. Data are represented as mean ± SD **(a, b)** or ± SEM **(e, f)**. Statistical significance was determined using two-way ANOVA with Tukey’s multiple comparisons test (\**p*<0.05, **p<0.01, \*\*\**p*<0.001).

Given the close association between behavioural deficits and nigrostriatal dopaminergic loss, we next examined whether mFcγRII deletion mitigated PFF-induced neurodegeneration. We assessed expression of tyrosine hydroxylase (TH) as a marker of catecholamine neurons [32] and dopaminergic neuron integrity. At 90 dpi, WT mice injected with PFF exhibited a marked reduction in TH⁺ fibre density within the ipsi dorsal striatum (Fig. 4c, e), accompanied by a significant loss of TH⁺ neuronal somata in the ipsi SNpc (Fig. 4d, f) relative to both Mono α-Syn-injected controls (Fig. 4c–f) and the 30 dpi cohort (Fig. S7). By contrast, mFcγRII^-/-^ mice showed comparable TH⁺ fibre densities and neuronal soma counts following either PFF or Mono α-Syn injection at both 30 dpi (Fig. S7) and 90 dpi (Fig. 4c-f). This suggests that mFcγRII deletion confers robust protection against both dopaminergic signalling deficits and the associated motor dysfunction in the PFF model.

### Prophylactic mAb-mediated hFcγRIIb blockade mitigates PFF-induced α-Syn Pathology

To determine whether FcγRIIb actively modulates the emergent stages of pathological α-Syn propagation *in-vivo*, and assess the therapeutic potential of targeting FcγRIIb, we evaluated whether prophylactic blockade of hFcγRIIb attenuates nascent pα-Syn formation in hFcγRIIb Tg mice [23]. Expression of the hFcγRIIb transgene, along with the absence of mFcγRII, was confirmed in the brain *via* RT-PCR, detecting human *FCGR2B* transcripts in naïve hFcγRIIb Tg mouse brain homogenates (Fig. 5a). Complementary immunohistochemistry in brain (Fig. 5b) and immunofluorescence in spleen (Fig. 5c) verified physiologically relevant expression, validating this model for translational interrogation of human-selective FcγRIIb antibodies.

**Figure 5.**
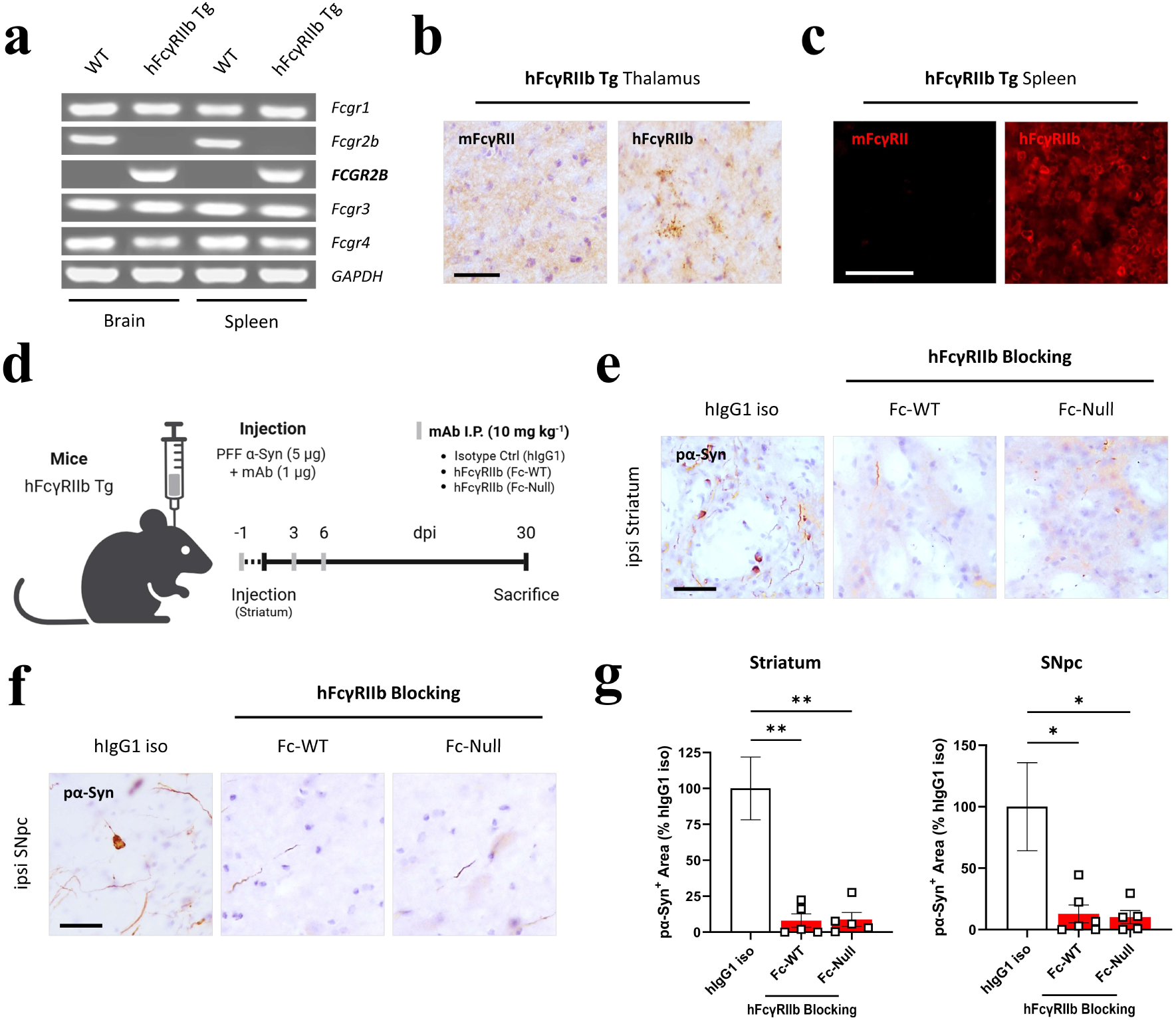
Acute-phase pα-Syn accumulation is reduced in hFcγRIIb Tg mice treated with hFcγRIIb blocking mAbs. **(a)** Agarose gel showing FcγR transcript expression in naïve mouse tissues. Total RNA from brain and spleen hFcγRIIb Tg mice was analysed by RT-PCR using primers mFcγRI (*Fcgr1*), mFcγRII (*Fcgr2b*), hFcγRIIb (*FCGR2B*), mFcγRIII (*Fcgr3*), mFcγRIV (*Fcgr4*), and *GAPDH*. WT mice transcripts serve as a comparator. **(b)** Representative images of immunohistochemical mFcγRII and hFcγRIIb staining in the thalamus from hFcγRIIb Tg mice. Sections were counterstained with haematoxylin. Scale bar = 50 μm. **(c)** Representative images of immunofluorescent mFcγRII and hFcγRIIb staining (red) in the spleen of hFcγRIIb Tg mice. Scale bar = 200 μm. **(d)** Schematic representation of *in-vivo* study design. **(e, f)** Representative images of immunohistochemical pα-Syn staining in **(e)** the ipsilateral (ipsi) striatum and **(f)** ipsi Substantia Nigra *pars compacta* (SNpc) of hFcγRIIb Tg mice injected with PFF α-Syn: co-administered with hIgG1 isotype (iso) control, hFcγRIIb blocking Fc-WT, or hFcγRIIb blocking Fc-Null mAbs at 30 dpi. Scale bars = 50 μm. **(g)** Quantification of pα-Syn^+^ immunoreactivity at 30 dpi in indicated brain regions and groups from **(e, f)** represented as a % of hIgG1 iso control. hIgG1 iso; *n* = 6, hFcγRIIb blocking Fc-WT; *n* = 6, hFcγRIIb blocking Fc-Null; *n* = 5 animals. Data are represented as mean ± SEM. Statistical significance was determined using one-way ANOVA with Tukey’s multiple comparisons test (\**p*<0.05, \*\**p*<0.01).

A mixed-sex cohort of adult hFcγRIIb Tg mice received intracranial co-administration of 5 µg PFF α-Syn together with 1 µg of a hFcγRIIb-specific mAb (*n* = 6 per group). Two hFcγRIIb blocking mAbs (hIgG1) were tested, incorporating either Fc-WT or Fc-Null (N297Q variant) Fc domains [23, 25]. Inclusion of both variants enabled resolution between effects arising solely from hFcγRIIb blockade from those potentially arising from Fc-mediated engagement of activating FcγRs on neighbouring immune cells. A hIgG1 isotype (iso) control mAb served as a comparator, and additional intraperitoneal dosing (10 mg kg^-1^) maintained circulating antibody levels (Fig. 5d).

At 30 dpi, immunohistochemical staining for pα-Syn revealed a near-complete abrogation of pα-Syn^+^ inclusions in the brains of hFcγRIIb blocking mAb treated mice. Quantitative assessment demonstrated a ∼90% decrease in pα-Syn^+^ burden within the ipsi striatum and SNpc of Fc-WT and Fc-Null mice relative to hIgG1 iso-treated controls (Fig. 5e-g). No ectopic pα-Syn immunoreactivity was detected elsewhere in the brain across all treatment groups, consistent with the restricted, early-stage distribution of pα-Syn mirrored in WT mice at 30 dpi (Fig. S3). These data suggest that prophylactic antibody-mediated hFcγRIIb blockade effectively suppresses early-stage pathologic α-Syn templating at the injection site and consequent propagation to anatomically connected regions like the SNpc.

## Discussion

The cell-to-cell transmission of pathological α-Syn aggregates constitutes a fundamental mechanism driving the progression of PD and related synucleinopathies [4, 7]. Previous studies have convincingly shown that aggregated α-Syn binds directly to FcγRIIb *in-vitro*, mediating uptake through lipid raft-dependent endocytic pathways [11, 21]. Moreover, receptor knockdown strategies undertaken during these investigations mitigated α-Syn uptake *in-vitro*. However, the contribution of FcγRIIb to the *in-vivo* dynamics governing propagation of α-Syn pathology has remained unexplored. Here, employing the established PFF model [27], we demonstrate that mFcγRII, the murine orthologue of hFcγRIIb, sharing 52% identity within the extracellular ligand-binding domain (https://www.uniprot.org/align/), serves as a critical determinant of α-Syn seeding competence and the trans-neuronal propagation of pathology across the mouse brain.

Genetic ablation of mFcγRII markedly reduced pα-Syn accumulation and distribution in response to PFF challenge at early (30 dpi) and intermediate (90 dpi) time points (Fig. 2). The attenuated pathological burden was accompanied by preservation of TH^+^ dopaminergic neurons within the SNpc and amelioration of motor deficits (Fig. 4), strongly implicating mFcγRII-mediated fibril uptake as an essential step in α-Syn-induced neurotoxicity. The absence of Mono α-Syn-induced pathology across genotypes confirms that the effects of α-Syn on mFcγRII are restricted to seeding-competent fibrillar α-Syn species [10, 11].

Other cell-surface receptors have been implicated in α-Syn uptake. For example, LAG3 deletion reduces neuronal internalisation, delays pathology *in-vivo*, and protects dopamine neurons [33], while APLP1 loss similarly diminishes propagation and neurotoxicity [34]. Other candidates, including the α3 subunit of Na⁺/K⁺ ATPase [35] and connexin32 [36], also remain under investigation. This indicates that α-Syn fibrils may engage a diverse repertoire of receptors, akin to Aβ, highlighting the potential for multiple, possibly overlapping uptake pathways. Nonetheless, the overt reduction in pα-Syn burden observed in mFcγRII^-/-^ mice suggests that FcγRIIb represents a mechanistically distinct and influential conduit for α-Syn propagation. While differences in experimental design preclude direct quantitative comparison with LAG3 knockout studies [33], our findings indicate that mFcγRII contributes substantially to the efficiency of α-Syn propagation and resulting neurotoxicity *in-vivo*.

The absence of mFcγRII was also associated with a markedly dampened microglial response to PFF. Although FcγRIIb is conventionally considered an inhibitory receptor that restrains immune activation [12, 16], its deletion did not exacerbate microgliosis in the model (Fig. 3). Compared to WT mice, mFcγRII^-/-^ mice exhibited insubstantial Iba1 and FcγRI expression at both 30 dpi and 90 dpi. This likely reflects the diminished α-Syn burden observed in these mice, consonant with evidence that microglial responses scale with extracellular aggregate burden [30]. These observations may also suggest that mFcγRII’s primary contribution to α-Syn pathology *in-vivo* is neuron-intrinsic, with reduced fibril internalisation limiting secondary glial activation. However, further investigation is warranted to dissect the acute *vs.* chronic immune consequences of mFcγRII deletion in the context of α-Syn-induced neuroinflammation and the role of microglia *in-vivo* [10].

To evaluate the translational potential of this axis, we implemented a prophylactic mAb blockade strategy using hFcγRIIb mAbs, which selectively antagonise hFcγRIIb with high specificity [23, 24]. Both the Fc-WT and Fc-Null variants successfully impaired acute α-Syn seeding competence in the striatum (site of injection), thereby disrupting downstream propagation towards the SNpc of hFcγRIIb Tg mice at 30 dpi (Fig. 5). The comparable efficacy of the Fc-WT and Fc-Null mAbs suggests that blockade is mediated predominantly *via* steric interference with the putative α-Syn fibril-binding interface on hFcγRIIb. Although the precise PFF-hFcγRIIb docking site has yet to be structurally resolved, it is thought to involve both extracellular Ig-like domains [11], and partial overlap of the Fc-binding groove could explain the observed efficacy of receptor blockade. By acting directly on hFcγRIIb, these antibodies exploit a mechanistically distinct pathway from conventional anti-α-Syn immunotherapies, offering a novel and perhaps complementary approach to suppress neuronal uptake and subsequent seeding of pathogenic α-Syn.

The mechanistic complementarity of Fc-WT and Fc-Null hFcγRIIb mAbs also highlights the translational versatility afforded by antibody engineering strategies to minimise off-target immune activation. For instance, the Fc-WT mAb can elicit B cell depletion *via* engagement with activating FcγRs, whereas the Fc-Null variant, incorporating an N297Q substitution, retains potent hFcγRIIb blockade without triggering this adverse effect [23, 24]. Such Fc modifications are increasingly exploited in therapeutic antibody development, though parameters like antibody half-life, neonatal Fc receptor interactions, and residual low-affinity FcγR binding must be carefully considered [14]. Importantly, the mAbs used in this study (Fc-WT [BI-1206] and Fc-Null [BI-1607]), are currently undergoing Phase I/IIa clinical trials in patients with haematological malignancies (Clinical Trial ID: <u>NCT03571568</u>) and solid cancers (Clinical Trial ID: <u>NCT04219254</u> and <u>NCT06784648</u>). This existing clinical development pathway is expected to facilitate their rapid translation into therapeutic strategies for neurodegenerative disease, pending further validation.

Several limitations of our study warrant consideration. First, while intracerebral antibody delivery affords proof-of-concept mechanistic dissection, it does not recapitulate the pharmacokinetics or bio-distribution of systemic administration, critical for clinical translation. Moreover, our prophylactic paradigm does not address therapeutic intervention at stages of established pathology, which would be essential to reflect the clinical context of patients with PD. Second, our initial study using mFcγRII^-/-^ mice was confined to male mice. Given the growing recognition of sex-specific immune responses in PD [37], future work must incorporate both sexes to assess generalisability. Third, although our data implicate FcγRIIb in early-mid stage α-Syn propagation, its role in later-stage neurodegeneration or α-Syn clearance remains to be elucidated.

In conclusion, our findings establish FcγRIIb as a pivotal mediator of α-Syn propagation *in-vivo*, acting as a neuron-intrinsic fibril uptake pathway that dictates the efficiency of early α-Syn seeding and modulating secondary glial activation. Targeted hFcγRIIb blockade using mAbs effectively disrupts acute pα-Syn deposition, representing a mechanistically precise, potentially disease-modifying strategy. When integrated with the existing clinical precedent for FcγRIIb-directed therapeutics in oncology [23, 38, 39], these data support the feasibility of repurposing or optimising receptor-specific interventions to arrest the progression of α-synucleinopathies.

## Acknowledgements

The authors thank the staff of the Biomedical Research Facility and the Biomedical Imaging Unit at the University of Southampton for their expert technical support. We also acknowledge additional support from colleagues at the University of Southampton, including Katrin Deinhardt, Cormac Spence, and Phil Williamson, for their valuable input throughout the study. This work was supported by a Wessex Medical Research PhD studentship awarded to A.R. and J.T. to support J.H. Additional funding was generously provided to J.H. by the Gerald Kerkut Charitable Trust.

## Methods

### Mice

WT C57BL/6 (Charles River Laboratories), mFcγRII^-/-^ [22], and hFcγRIIb ^+/-^ Tg mice [23] were bred and maintained at the Biomedical Research Facility (University of Southampton). Additionally, hFcγRIIb Tg mice were intercrossed with mFcγRII^-/-^ mice to eliminate endogenous mFcγRII expression. Genotypes were confirmed prior to experimentation using flow cytometry of peripheral blood and PCR-based amplification of the hFcγRIIb transgene from ear tip biopsies (data not shown). All experiments were performed in accordance with UK Home Office regulations (under PIL I84150147 and PPL PP6936815) and approved by the University of Southampton’s Animal Welfare and Ethical Review Body. Mice were maintained under standard housing conditions (21 ± 2°C, 45-60% humidity, 12-hour light/dark cycle) with environmental enrichment and *ad libitum* access to irradiated RM1 (E) diet and water. Mice were age-matched within experiments and groups assigned based on genotype and/or transgene expression [23].

### Purification of Recombinant α-Syn and Preparation of PFF α-Syn

Recombinant human WT α-Syn was expressed in *Escherichia coli* BL21 RIPL pLacI cells (Agilent) transformed with a pET-α-Syn plasmid. The plasmid was kindly provided by Phil Williamson and generated by cloning α-Syn DNA into a pET vector (Merck). Protein expression was induced with isopropyl β-D-1-thiogalactopyranoside and purified using Q-Sepharose anion exchange chromatography and gel filtration under denaturing conditions. Refolding was performed in an oxido-reduction buffer, followed by dialysis and re-purification on Q-Sepharose. Endotoxin removal was performed using the 0.1% Triton X-114 method as described by Reichelt et al. [40]. Purified protein was buffer-exchanged into PBS, sterile-filtered, and stored at -80°C. Endotoxin levels, determined using the Endosafe^®^ NexGen-PTS^TM^ system (Charles River Laboratories), were < 1 EU mg⁻¹, and protein purity exceeded 95% as confirmed by Coomassie-stained SDS-PAGE. Mono α-Syn (5 mg mL⁻¹) was aggregated at 37°C under continuous agitation (1000 rpm) for 7 days. The resulting fibrils were fragmented into PFF particles (< 50 nm) *via* sonication in a cup-horn ultrasonic water bath (Fisherbrand™ Model 705). PFF formation was validated per preparation by transmission electron microscopy and Thioflavin-T Fluorimetry.

### *In-Vitro* Experiments

#### Mouse Primary Neuron Experiment

Cortical neurons were isolated from E15.5 WT or mFcγRII^-/-^C57BL/6 mouse embryos. Following dissection, tissue was enzymatically dissociated in 0.05% trypsin at 37°C for 5 minutes, followed by gentle trituration to obtain a single-cell suspension. Cells were plated on poly-D-lysine-coated glass coverslips in 24-well plates and maintained in Neurobasal™ medium supplemented with 2% (v/v) B27™ and 0.5 mM GlutaMAX™ (Gibco) at 37°C in 5% CO2. After 10 days, neurons were exposed to 1 µg PFF α-Syn or BSA (Sigma) per well and maintained for an additional 14 days. Each experiment was conducted in triplicate across four independent neuronal preparations.

#### Immunocytochemistry

Cells were fixed in 4% paraformaldehyde (PFA) in 0.1 M phosphate buffer (PB) at RT for 20 minutes and rinsed with PBS. Non-specific binding was blocked for 90 minutes using 10% normal goat serum (NGS) in blocking buffer (PBS containing 0.005% Triton X-100 and 0.05% thimerosal, pH 7.4), followed by overnight incubation at 4°C in blocking buffer with 2% NGS and purified mouse anti-pSer129-α-Syn (BioLegend^®^). The following day, a biotinylated goat anti-mouse secondary antibody (Vector Laboratories) was applied for 2 hours at RT in blocking buffer with 2% NGS. Immunoreactivity was detected using streptavidin–horseradish peroxidase (ThermoFisher) and visualised with 0.05% 3, 3’-Diamninobenzindene (DAB) with 0.015% hydrogen peroxide. Coverslips were rinsed with PBS, detached, and mounted onto microscope slides with Mowiol^®^ (Sigma).

### *In-Vivo* Experiments

#### Genotype Experiment

All surgical procedures were carried out under aseptic conditions. Adult male WT and mFcγRII^-/-^ C57BL/6 mice (8 weeks old, *n* = 3-4 per group) were anaesthetised with isoflurane (inhaled) and positioned in a stereotaxic frame (KOPF Instruments). A Hamilton™ 10 µL syringe (GASTIGHT™ 1700 series) was used to unilaterally inject 5 µg PFF or Mono α-Syn (1 µL of 5 µg µL⁻¹) at 0.4 µL min⁻¹ at the following coordinates from Bregma: AP +0.4 mm, ML +2.0 mm, DV -3.0 mm. The syringe was left in place for ≥ 5 minutes post-injection to facilitate diffusion before withdrawal. The incision was sutured and post-surgical care provided. Mice were euthanized at 30 or 90 dpi for immunohistochemical analysis.

#### hFcγRIIb mAb Blockade Experiment

Adult hFcγRIIb Tg mice (12-16 weeks old, *n* = 6 per group consisting of 4 males and 2 females) underwent unilateral stereotaxic surgery as described above. 5 μg PFF α-Syn (1 μL of 5 μg μL⁻¹) was co-injected with an hIgG1 hFcγRIIb blocking mAb: Fc-WT (BI-1206) or Fc-Null (BI-1607), or an hIgG1 iso control mAb (cetuximab). Antibodies were pre-mixed with PFF α-Syn prior to inoculation, with a total antibody dose of 1 μg per mouse. In addition, antibodies were administered intraperitoneally (I.P.) at a dosage of 10 mg kg⁻¹ to ensure sustained peripheral antibody load. I.P. injections were performed one day prior to surgery and again at 3 and 6 days post-surgery. One mouse from the Fc-Null group was culled early due to ill health following the surgical procedure. No other acute detrimental effect was observed following administration of the antibody in any of the other mice. Mice were euthanized 30 dpi for immunohistochemistry.

### Behavioural Testing

Mice from the genotype study (90 dpi cohort) underwent regular behavioural testing at monthly intervals prior to sacrifice. Testing occurred 4 times total: pre-surgery, 1, 2, and 3 months post-surgery. Mice were habituated to the procedure room and the investigator remained blind to experimental groups throughout.

#### Wire Hang

The wire hang test of grip strength and motor coordination evaluates neuromuscular function in murine models [41]. A mouse was positioned on a 35 cm long, 2 mm diameter wire suspended 30 cm above a padded surface. The latency to fall off the wire was recorded, and averaged across three trials with a maximum duration of 30 seconds per trial.

#### Pole Test

The pole test assesses motor coordination and balance in mice [41]. A mouse was placed near the top of a 60 cm long, 1 cm diameter vertical pole. The time taken to turn and descend to the base was measured over three trials, with a maximum duration of 120 seconds per trial.

#### Rotarod Test

The rotarod test is used to evaluate motor learning, coordination, and balance [42]. Prior to baseline assessments, mice were acclimated to the rotarod apparatus (Ugo Basile) for three 5 minute trials at 10 revolutions per minute over two consecutive days. On the test day, mice were placed on the accelerating rotarod (ranging from 4 to 40 rpm over 300 seconds), and the latency to fall was recorded for a maximum of 5 minutes. Each mouse received three consecutive trials and the mean latency to fall was used for analysis.

#### Open Field Test

The open field test is a standardised assessment of locomotor activity and anxiety-related behaviour in rodents [41]. Mice were habituated to the open field arena for 10 minutes across two consecutive days prior to baseline assessments. On the test day, subjects were placed in the centre of a 76 cm × 76 cm × 50 cm square arena for 10 minutes while locomotor activity was monitored using an automated infrared beam system (EthoVision). The initial 2 minutes of each session were excluded from analysis to account for initial excitability.

### Tissue Collection

Mice were terminally anesthetised (Avertin^®^), and transcardially perfused with ice-cold heparinised saline (5 units mL^-1^), followed by 4% PFA in 0.1 M PB. Tissues were post-fixed in 4% PFA at 4°C for 4 hours. For cryoprotection, brains were transferred to 30% sucrose in 0.1 M PB prior to sectioning. Coronal sections (40 μm) were obtained using a CM1860 UV cryostat (Leica) in 12-series and stored in antifreeze solution (30% ethylene glycol, 30% glycerol, 30% ddH2O, 10% 0.2 M PB) at -20°C until required. An additional cohort of naïve WT, mFcγRII^-/-^, and hFcγRIIb Tg mice (8-12 weeks old) were sacrificed for RT-PCR analysis of brain and spleen tissue. These mice were terminally anesthetised (Avertin^®^) and transcardially perfused with ice-cold heparinised saline. Brain tissues were snap-frozen in liquid nitrogen and stored at -80°C prior to homogenisation and RNA isolation.

### Immunohistochemistry

Immunohistochemistry was performed on 40 μm-thick free-floating coronal brain sections. Endogenous peroxidase activity was quenched (10% CH3OH, 3% H2O2 in ddH2O) for 10 minutes prior to blocking with 3% NGS in Tris-Buffered Saline (TBS) containing 0.2% Triton X-100 (TBS-T, pH 7.4) for 1 hour, followed by overnight incubation at RT with primary antibodies (Table S1) in 1% NGS in TBS-T. The following day, sections were incubated with biotin-conjugated secondary antibodies (Table S1) in TBS for 2 hours, followed by an avidin-biotin complex amplification step (Vector Laboratories) for 90 minutes. Immunoreactivity was visualised using DAB peroxidase substrate. Chromogen development was monitored, and the reaction was halted using several Tris-NaCl solution and TBS washes. Sections were mounted on gelatine-coated glass slides and allowed to air dry overnight. Mayer’s haematoxylin (Sigma) was applied as a counterstain, and sections were sequentially dehydrated in a graded series of ethanol solutions (70%, 95%, and 100%), cleared in xylene, and cover-slipped with DPX (Sigma). The mFcγRII and hFcγRIIb stains (Fig. 1b, Fig. 5b) underwent an additional heat-induced antigen retrieval step in Tris-EDTA buffer (10 mM Tris, 1 mM EDTA, 0.05% Tween-20, pH 9.0) for 10 minutes at 90°C prior to quenching.

### Immunofluorescence

Immunofluorescent IHC was performed on 40 μm-thick free-floating coronal brain or spleen sections. Blocking was with 3% NGS in TBS-T for 1 hour, followed by overnight incubation at RT with primary antibodies (Table S1) in 1% NGS in TBS-T. Immunoreactivity was revealed using appropriate fluorescent secondary antibodies (Table S1) conjugated to Alexa-fluor 488 or 568 (Invitrogen) in TBS for 2 hours. Nuclei were counterstained with 4′, 6-diamidino-2-phenylindole (DAPI; Merck) for 5 minutes. Stained sections were mounted on gelatin-coated glass slides and allowed to air dry overnight away from direct light. Slides were cover-slipped using Mowiol^®^ (Sigma).

### Reverse Transcription PCR

Total RNA was extracted from snap-frozen brain or spleen tissue using the RNeasy Mini Kit (Qiagen) according to the manufacturer’s instructions. Complementary DNA (cDNA) was synthesised from 400 ng of RNA using the TaqMan Reverse Transcription Kit (Applied Biosystems^TM^, Life Technologies). PCR was performed using the StepOnePlus™ Real-Time PCR System (Applied Biosystems^TM^, Thermo Fisher Scientific) with gene-specific primers (Table S1). All reactions were performed in technical duplicate. RT-PCR products were resolved by agarose gel electrophoresis and visualised using GelRed™ nucleic acid stain (Biotium) on a LI-COR Odyssey Fc imaging system to confirm amplicon size and specificity.

### Image Acquisition and Analysis

Quantification of immunoreactivity was performed using unbiased, design-based stereological approaches. For primary neuronal cultures, five randomly selected fields of view were imaged per condition from each of three replicate coverslips (15 images in total). For tissue derived from *in-vivo* experiments, three serial 40 µm coronal sections per animal encompassing the relevant regions of interest (ROIs) were selected for analysis. Within each section, images were acquired using a 40× objective from five randomly chosen fields of view per ROI (15 images per ROI per mouse). Image acquisition parameters were kept constant across groups. Quantification of staining was performed using FIJI (ImageJ v1.53h), measuring the percentage area of positive staining per field of view. Distribution maps of pα-Syn⁺ immunoreactivity were generated in FIJI to depict the average spatial distribution of Lewy-like pathology within each experimental group. TH⁺ neurons in the SNpc were manually counted from images captured at 20× magnification and corrected for total SNpc volume using the Cavalieri principle [43]. Optical density of TH immunoreactivity in the dorsal striatum was measured in FIJI from images acquired at 10× magnification. All analyses were performed by an investigator blinded to experimental groups.

### Statistical Analysis

Data analysis was conducted and visualised with Prism™ v.10 (GraphPad), presented as bar charts with ± SEM or SD error bars and all data points. Normality of the data was assessed using the Shapiro-Wilk test where *p*>0.05 assumed normal distribution. Statistical analysis and *post hoc* comparisons were completed as indicated in figure legends. A significance threshold of *p*<0.05 was adopted for all statistical tests.

## Supplementary Materials

**Supplementary Table 1.**
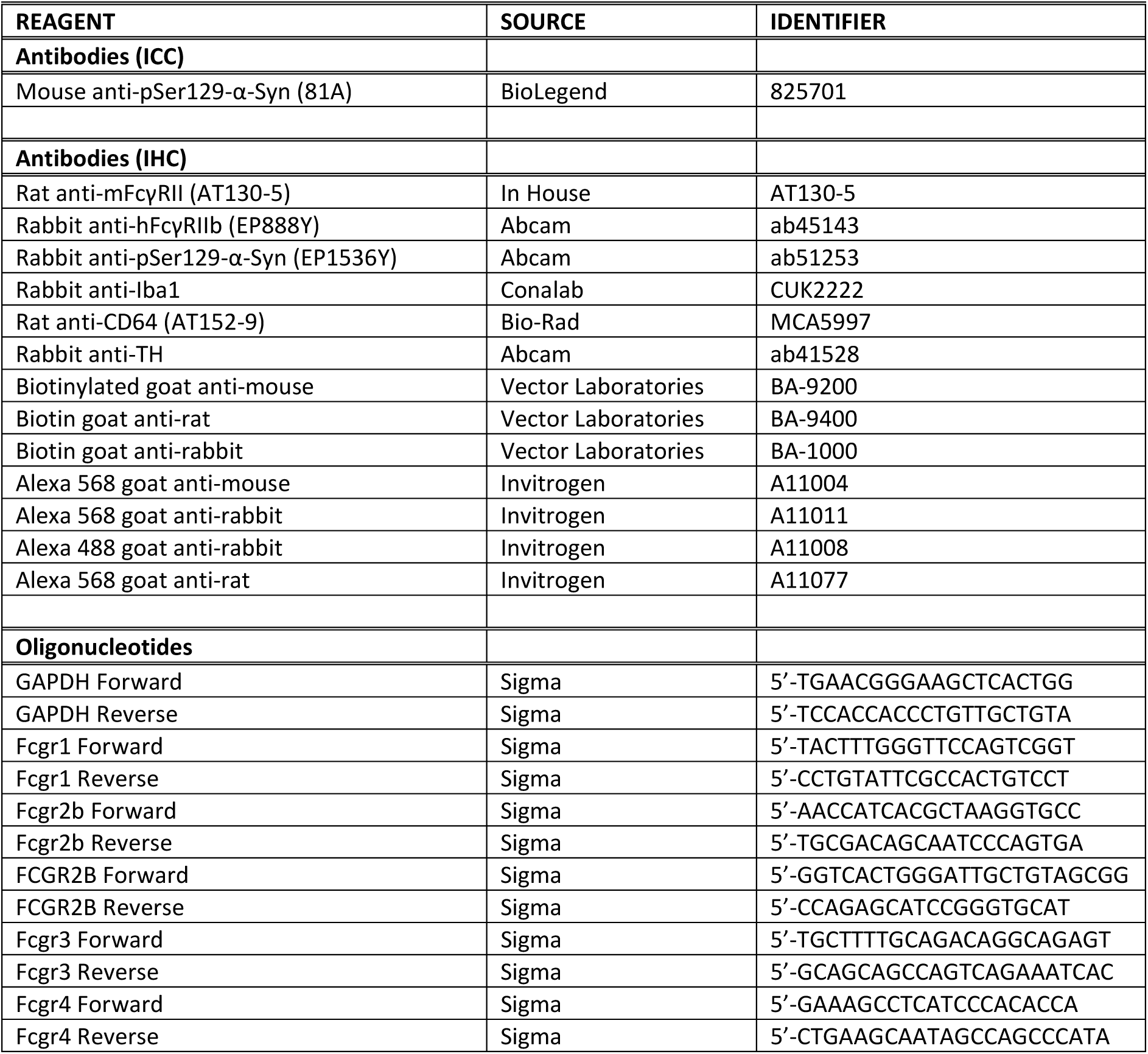
List of Antibodies and Oligonucleotides.

**Figure S1.**
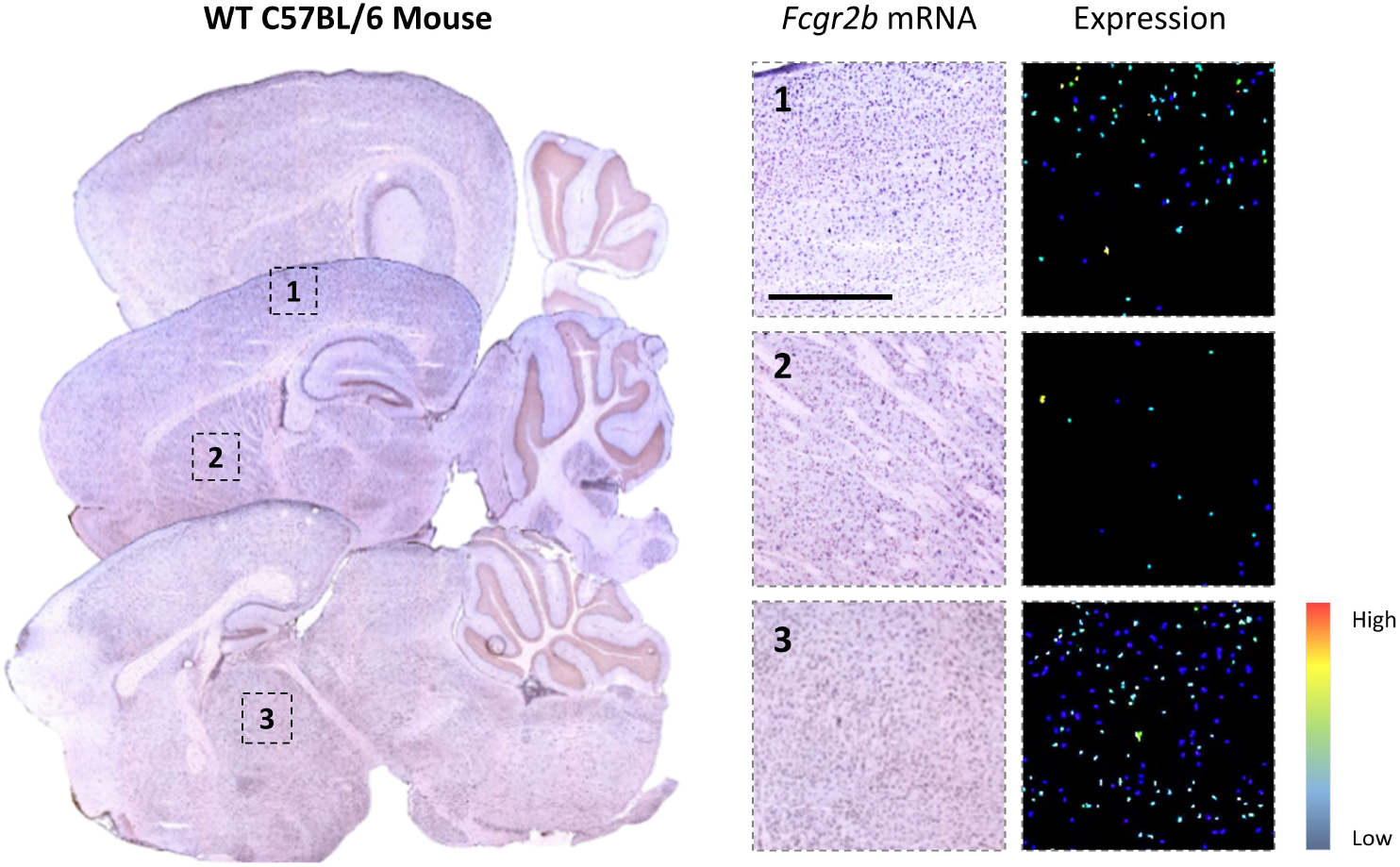
Expression pattern of *Fcgr2b* in the WT mouse brain from the Allen Mouse Brain Atlas. *In-situ* hybridisation and corresponding expression heat map illustrating *Fcgr2b* mRNA distribution in sagittal sections of the adult WT C57BL/6 mouse brain (postnatal day 56). Heat-map rendering highlights regional differences in expression, with enlarged panels depicting representative areas of interest. Scale bar = 500 μm. Data source: Allen Brain Atlas, © 2015 Allen Institute for Brain Science (https://mouse.brain-map.org/).

**Figure S2.**
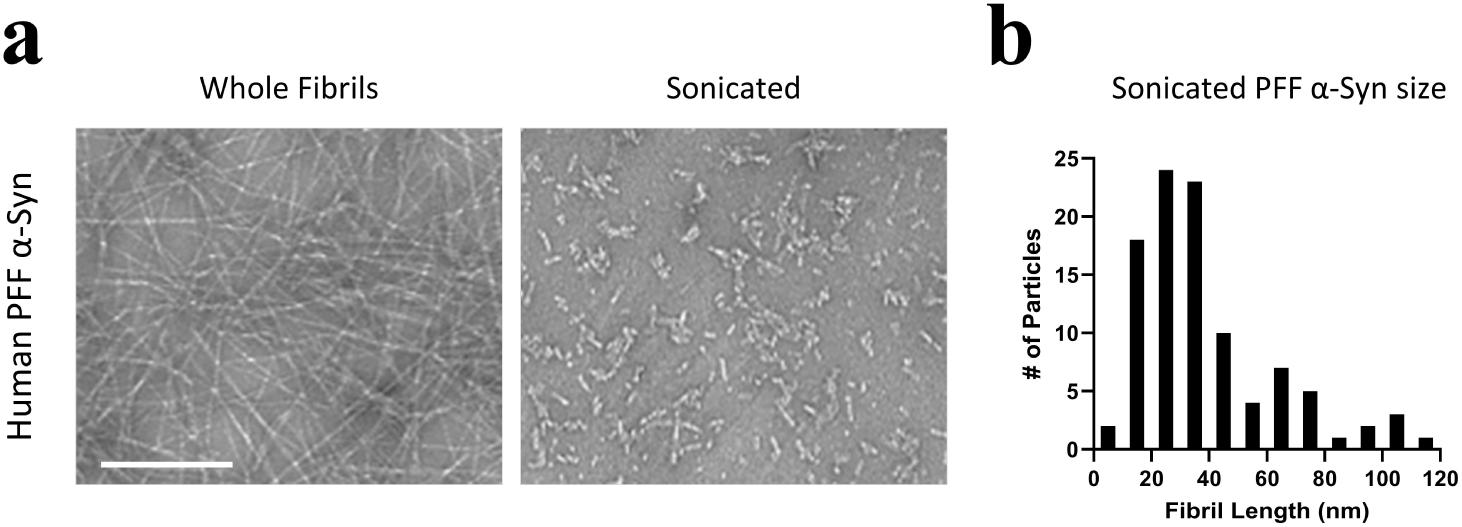
Preparation of recombinant human PFF α-Syn. **(a)** Representative transmission electron microscopy images of whole human PFF α-Syn and sonicated PFF α-Syn seeds. Scale bar = 200 nm. **(b)** Graph showing the distribution of PFF seed length post-sonication. Mean length = 34.3 nm ± 2.37 (*n* = 100).

**Figure S3.**
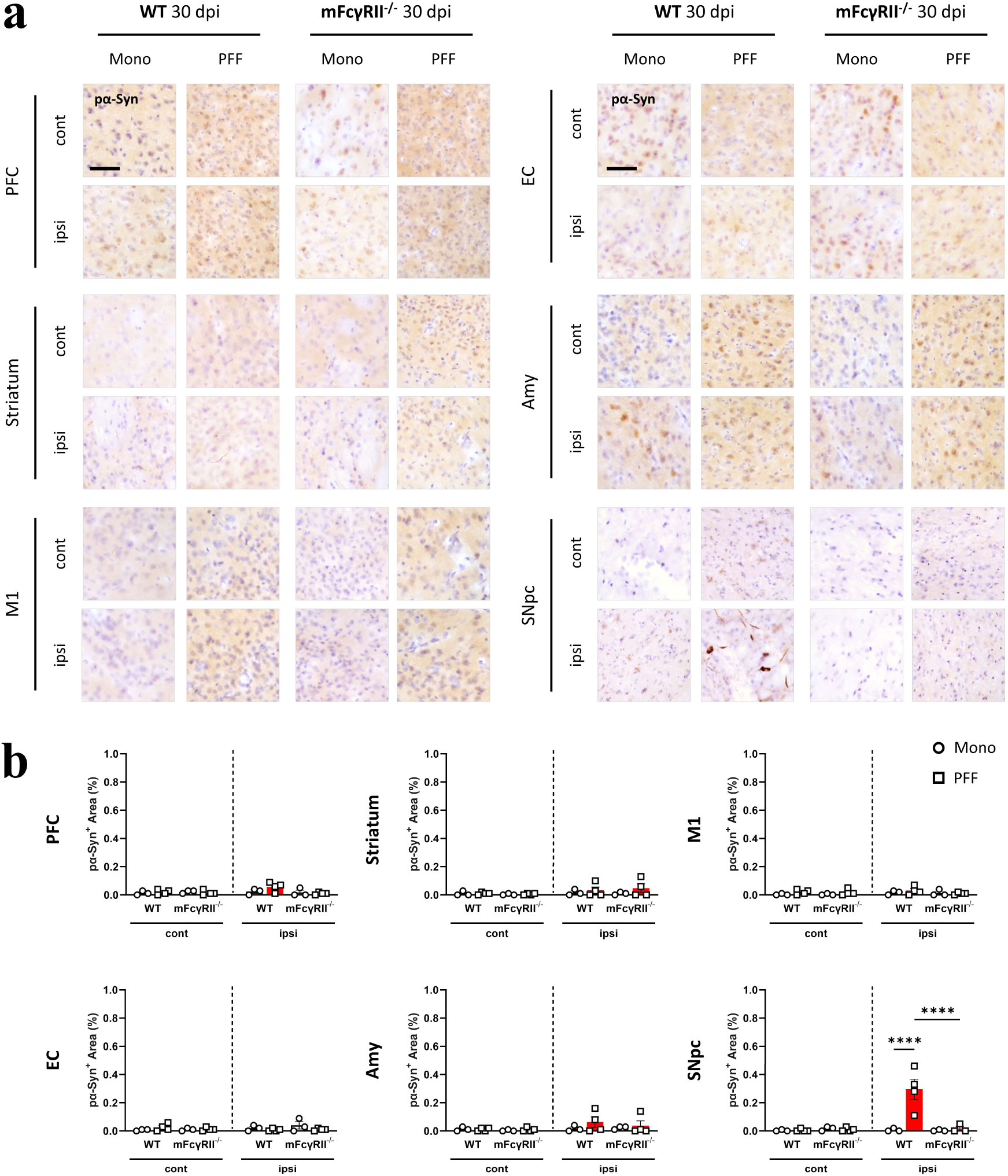
pα-Syn accumulation in WT and mFcγRII^-/-^ mice at 30 dpi. **(a)** Representative images of immunohistochemical pα-Syn staining in contralateral (cont) and ipsilateral (ipsi) hemispheres of WT or mFcγRII^-/-^ mice injected with PFF or Mono α-Syn at 30 dpi. PFC; prefrontal cortex, M1; primary motor cortex, EC; entorhinal cortex, Amy; amygdala, SNpc; substantia nigra *pars compacta*. Sections were counterstained with haematoxylin. Scale bar = 50 μm. **(b)** Quantification of pα-Syn^+^ immunoreactivity (% area) at 90 dpi in indicated brain regions and groups from **(a)**. WT Mono; *n* = 3, mFcγRII^-/-^ Mono; *n* = 3, WT PFF; *n* = 4, mFcγRII^-/-^ PFF; *n* = 4 animals. Data are mean ± SEM. Statistical significance was determined using two-way ANOVA with Tukey’s multiple comparisons test (\*\*\*\**p*<0.0001).

**Figure S4.**
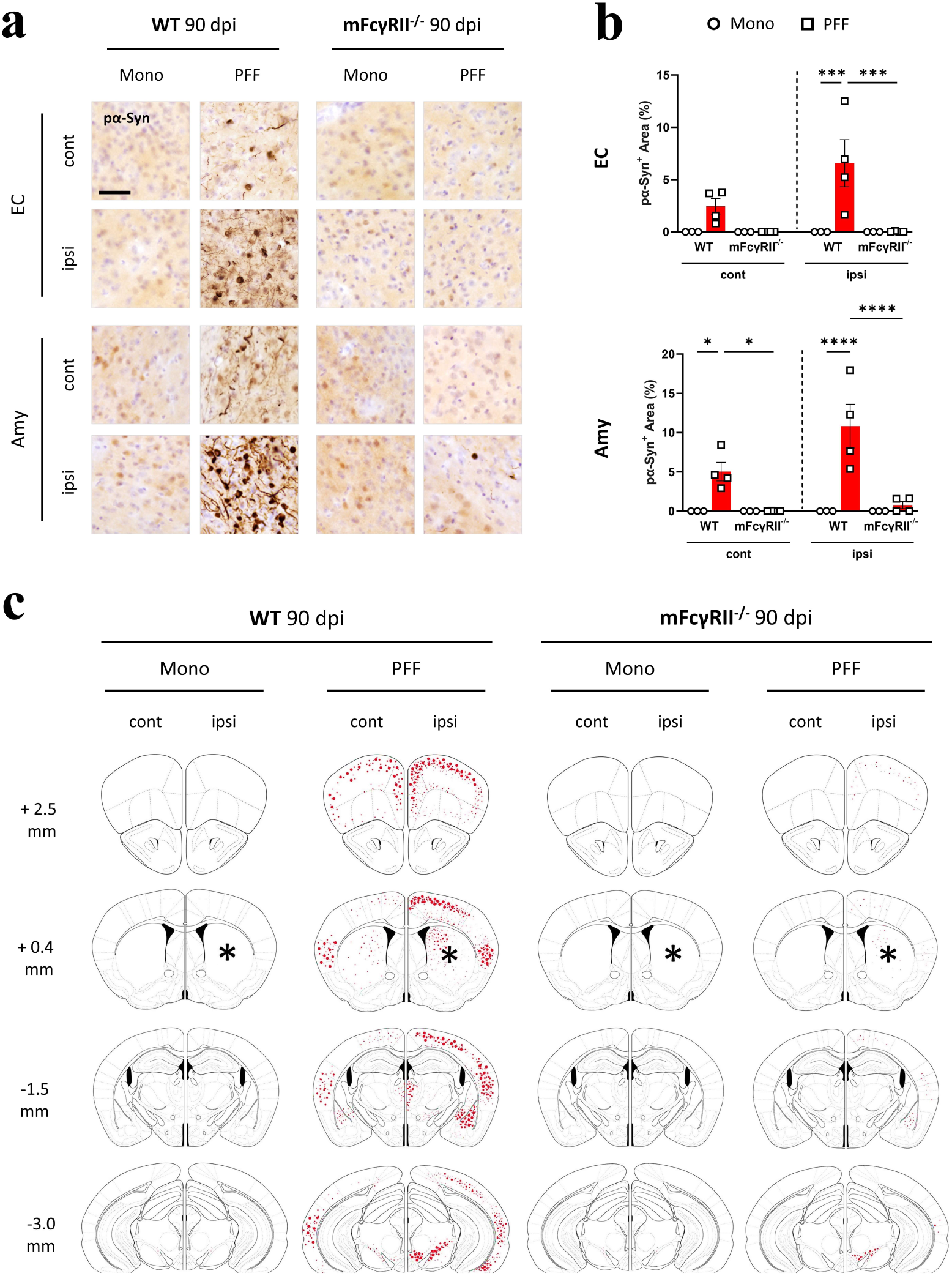
Regional distribution of pα-Syn in WT and mFcγRII^-/-^ mice at 90 dpi (additional brain regions). **(a)** Representative images of immunohistochemical pα-Syn staining in contralateral (cont) and ipsilateral (ipsi) hemispheres of WT or mFcγRII^-/-^ mice injected with PFF or Mono α-Syn at 90 dpi. EC; entorhinal cortex, Amy; amygdala. Sections were counterstained with haematoxylin. Scale bar = 50 μm. **(b)** Quantification of pα-Syn^+^ immunoreactivity (% area) at 90 dpi in indicated brain regions and groups from **(a)**. WT Mono; *n* = 3, mFcγRII^-/-^Mono; *n* = 3, WT PFF; *n* = 4, mFcγRII^-/-^ PFF; *n* = 4 animals. Data are mean ± SEM. Statistical significance was determined using two-way ANOVA with Tukey’s multiple comparisons test (\**p*<0.05, \*\*\**p*<0.001, \*\*\*\**p*<0.0001). **(c)** Distribution map of pα-Syn^+^ immunoreactivity (represented as red dots) across the whole brain at 90 dpi. Experimental groups are indicated alongside both contralateral (cont) and ipsilateral (ipsi) hemispheres. Asterisks denote the injection site (ipsi striatum), and adjacent numbers represent distance from Bregma in the anterior-posterior plane in mm.

**Figure S5.**
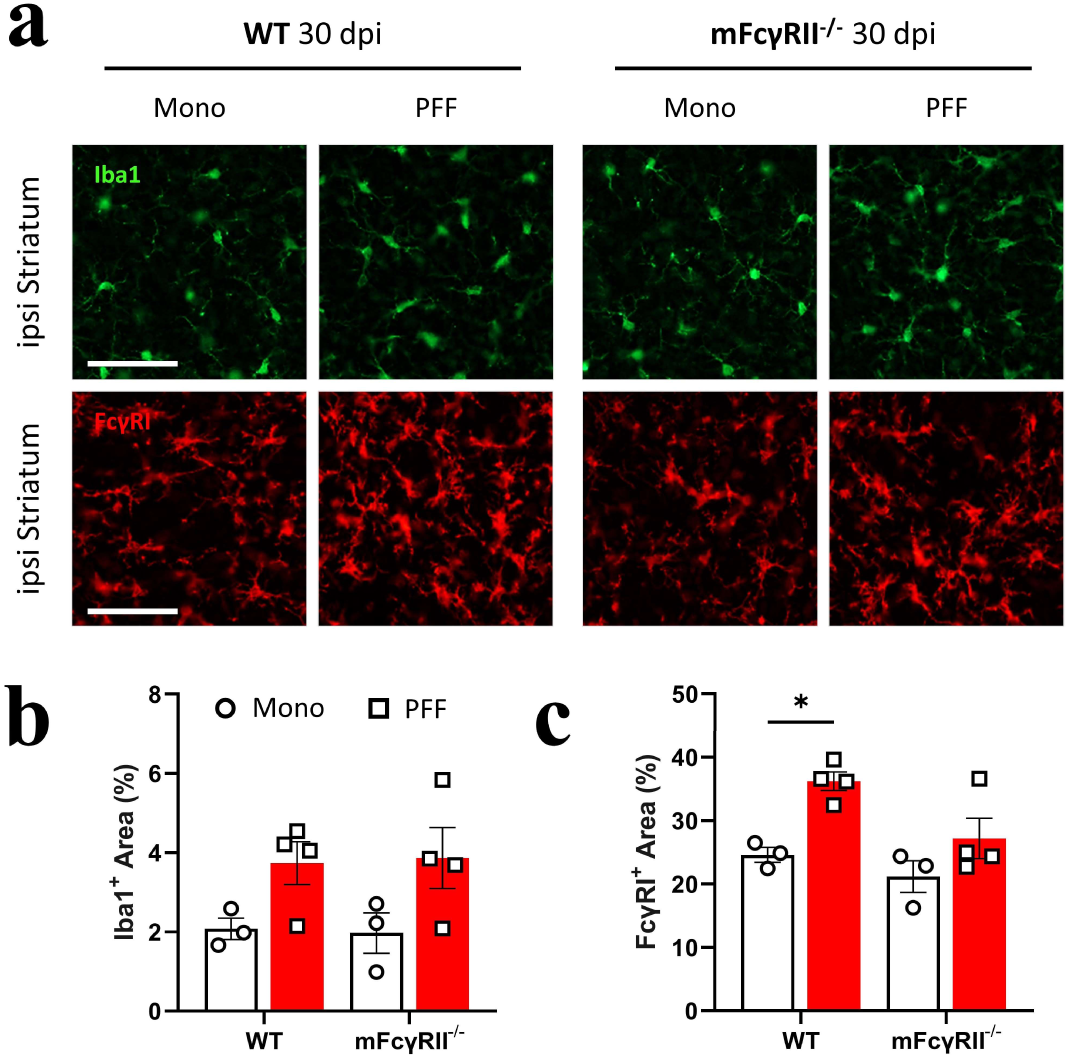
Microglial reactivity in the striatum of WT and mFcγRII^-/-^ mice at 30 dpi. **(a)** Representative images of immunofluorescent Iba1 (green) and FcγRI (red) in the ipsilateral (ipsi) striatum of WT and mFcγRII^-/-^ mice injected with PFF or Mono α-Syn at 30 dpi. Scale bars = 50 μm. **(b, c)** Quantification of **(b)** Iba1^+^ and **(C)** FcγRI^+^ immunoreactivity (% area) at 30 dpi in the ipsi striatum. WT Mono; *n* = 3, mFcγRII^-/-^ Mono; *n* = 3, WT PFF; *n* = 4, mFcγRII^-/-^ PFF; *n* = 4 animals. Data are mean ± SEM. Statistical significance was determined using two-way ANOVA with Tukey’s multiple comparisons test (\**p*<0.05).

**Figure S6.**
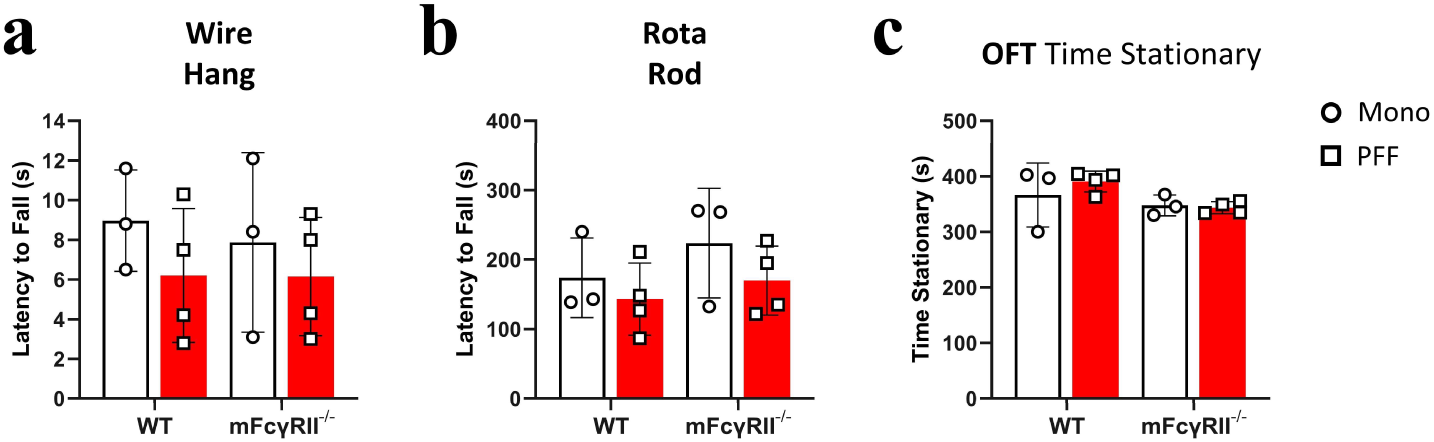
Additional behavioural assessments in WT and mFcγRII^-/-^ mice at 90 dpi. **(a)** Wire hang test assessing neuromuscular strength and coordination. Each point represents the latency to fall from the apparatus averaged over 3 trials per animal. **(b)** Rotarod test evaluating motor performance and balance. Each point represents the latency to fall from the apparatus averaged over 3 trials per animal. **(c)** Open-field test (OFT) analysis showing total time spent stationary. WT Mono, *n* = 3; mFcγRII^-/-^ Mono, *n* = 3; WT PFF, *n* = 4; mFcγRII^-/-^ PFF, *n* = 4 animals. Data are presented as mean ± SD.

**Figure S7.**
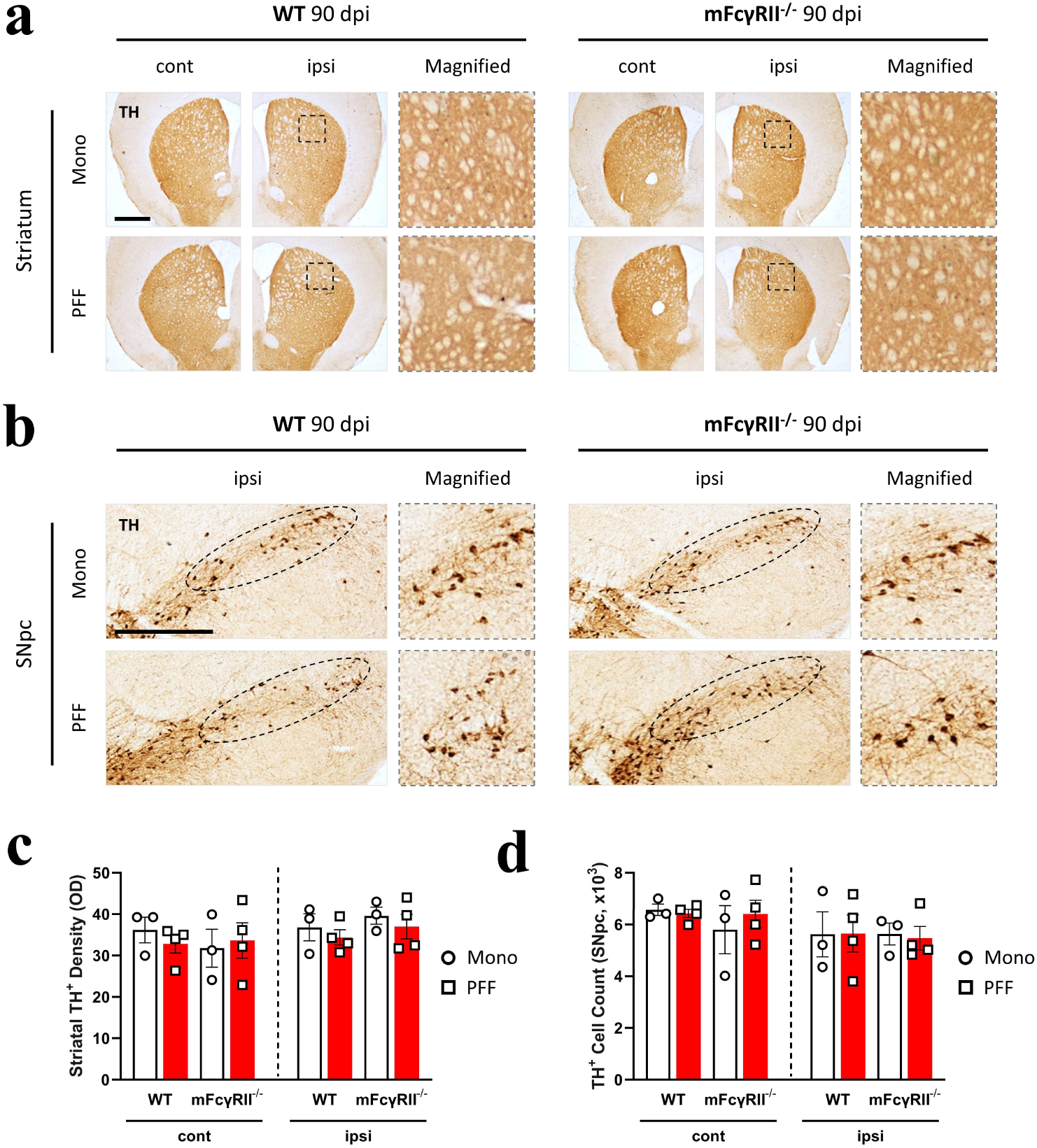
TH expression in WT and mFcγRII^-/-^ mice at 30 dpi. (a, **b)** Representative images of immunohistochemical TH staining in **(a)** the contralateral (cont) and ipsilateral (ipsi) striatum and **(b)** the ipsi Substantia Nigra *pars compacta* (SNpc) of WT and mFcγRII^-/-^ mice injected with PFF or Mono α-Syn at 30 dpi. Magnified images are shown adjacent to each panel. Scale bars = 1 mm **(a)** and 200 μm **(b)**. **(c)** Quantification of TH^+^ optical density (OD) in the dorsal striatum and **(d)** TH^+^ cells in the SNpc represented as a number normalised to the total SNpc volume *via* the Cavalieri estimate method. WT Mono; *n* = 3, mFcγRII^-/-^ Mono; *n* = 3, WT PFF; *n* = 4, mFcγRII^-/-^ PFF; *n* = 4 animals. Data are represented as mean ± SEM.

